# Promoter-Enhancer Interactions Identified from Hi-C Data using Probabilistic Models and Hierarchical Topological Domains

**DOI:** 10.1101/101220

**Authors:** Gil Ron, Dror Moran, Tommy Kaplan

## Abstract

Proximity-ligation methods as Hi-C allow us to map physical DNA-DNA interactions along the genome, and reveal its organization in topologically associating domains (TADs). As Hi-C data accumulate, computational methods were developed for identifying domain borders in multiple cell types and organisms.

Here, we present PSYCHIC, a computational approach for analyzing Hi-C data and identifying Promoter-Enhancer interactions. We use a unified probabilistic model to segment the genome into domains, which we merge hierarchically and fit the Hi-C interaction map with a local background model. This allows us to estimate the expected number of interactions for every DNA-DNA pair, thus identifying over-represented interactions across the genome.

By analyzing published Hi-C data in human and mouse, we identified hundreds of thousands of putative enhancers and their target genes in multiple cell types, and compiled an extensive genome-wide catalog of gene regulation in human and mouse.

## Introduction

One of the key mechanisms of gene regulation in eukaryotes involves enhancer-promoter interactions, where distal regulatory regions along the DNA (enhancers) come in close physical proximity to their target promoters, to further activate transcription. The human genome is estimated to contain hundreds of thousands of enhancers, often with multiple enhancers regulating a single gene. These act in a tissue specific manner and could be found up to 1Mb away from their target genes (Fraser and Bickmore 2007, Visel et al. 2009, Van Steensel and Dekker 2010, Bickmore and van Steensel 2013, Dekker and Mirny 2016, Rowley and Corces 2016). The importance of enhancers for gene regulation is further emphasized by a growing body of works that link genetic variation in enhancer sequences to human diseases (Lettice et al. 2003, Claussnitzer et al. 2015, Lupiáñez et al. 2015, Achinger-Kawecka and Clark 2016, Franke et al. 2016).

Nonetheless, we still lack a deep understanding of how enhancers work molecularly, how their tissue specificity is encoded in their DNA sequence, and above all how they recognize and physically interact with their target genes.

In recent years, high-throughput molecular methods have been developed to study the three-dimensional organization of the genome, and its relation to various functions. For example, proximity ligation methods such as 4C, ChIA-PET and Hi-C quantify the frequency of DNA-DNA interactions in living cells and map the 3D organization of the genome in high resolution (Simonis et al. 2006, Lieberman-Aiden et al. 2009, Handoko et al. 2011, Jin et al. 2013, Kieffer-Kwon et al. 2013, Rao et al. 2014, Fraser et al. 2015, Lajoie et al. 2015, Mifsud et al. 2015). To date, Hi-C experiments were performed in a variety of organisms and cellular conditions, including many cell types and tissues.

While the genomic resolution of these data is often low, varying from few Kbs to 40Kb blocks, they were mainly used to identify and delineate topologically associating domains (TADs). These are continuous regions (hundreds of Kb to few Mbs) that were shown to be folded upon themselves into local compartments and facilitate high number of local (cis) DNA-DNA interactions (Dixon et al. 2012, Nora et al. 2012, de Laat and Duboule 2013, Rao et al. 2014).

In recent years, topological domains were studied extensively, and were shown to be related to replication domains (Pope et al. 2014, Dileep et al. 2015), to be largely conserved across evolution, and to play a crucial role in chromosome function (Ryba et al. 2010, Dixon et al. 2012, Gómez-Marín et al. 2015, Jager et al. 2015, Vietri Rudan et al. 2015, Taberlay et al. 2016).

TADs also play a key role in gene regulation, as they define the regulatory scope of enhancers. The domains boundaries were shown to act as regulatory “insulators” that prevent targeting genes outside of the enhancer domain (Doyle et al. 2014, Symmons et al. 2014). Disruptions of the chromosomal structure, either in human genetic disorders or by artificially deleting boundary elements (e.g. using CRISPR-Cas9), were shown to be associated with enhancer mis-regulation and aberrant gene expression (Zhang et al. 2013, Lupiáñez et al. 2015, Achinger-Kawecka and Clark 2016, Blinka et al. 2016, Franke et al. 2016, Fulco et al. 2016). While we still lack a deep understanding of the exact mechanisms by which topological domains are defined and maintained, TAD borders seem be enriched for highly transcribed genes (Dixon et al. 2012), as well as CTCF and cohesin binding sites (Demare et al. 2013, Seitan et al. 2013, Ong and Corces 2014, Zuin et al. 2014, Ing-Simmons et al. 2015, Nichols and Corces 2015, Tang et al. 2015, Vietri Rudan et al. 2015, Fudenberg et al. 2016).

As more and more 3D data accumulate, in a multitude of tissues and cellular conditions, algorithms were developed to analyze Hi-C data and partition the genome into a set of topological domains (Dixon et al. 2012, Ay et al. 2014, Lévy-Leduc et al. 2014, Fraser et al. 2015, Lajoie et al. 2015,

Adhikari et al. 2016, Chen et al. 2016, Xu et al. 2016). Most notable is the statistical method by Dixon et al (2012), which scans the genome by analyzing the set of DNA-DNA interactions for every locus, and identifies transitions from loci with mostly backward interactions to adjacent loci with mostly forward interactions. While this method is generally fast and robust, it is inherently biased towards short-range interactions that form the vast majority of DNA-DNA interactions. This method also ignores a visible feature of Hi-C maps - the hierarchal structure of sub-domains organized into larger domains (Fraser et al. 2015).

Here, we present PSYCHIC (Fig 1) - a three step modular algorithm to identify promoter-enhancer interactions. Briefly, we use a unified probabilistic model and a Dynamic Programming algorithm to find an optimal segmentation of each chromosome into topological domains; we next iteratively merge neighboring domains into hierarchical structures; and finally we fit each domain using a local background model. This allows us to identify over-represented DNA-DNA pairs, including enhancers and their target genes. We have analyzed Hi-C data from 15 conditions and cell types in mouse and human (Dixon et al. 2012, Rao et al. 2014, Fraser et al. 2015), and identified hundreds of thousands of over-represented interactions. This comprehensive genome-wide tissue-specific database of putative interactions between enhancers and their target genes would be of great interest to the scientific community.

**Figure 1.**
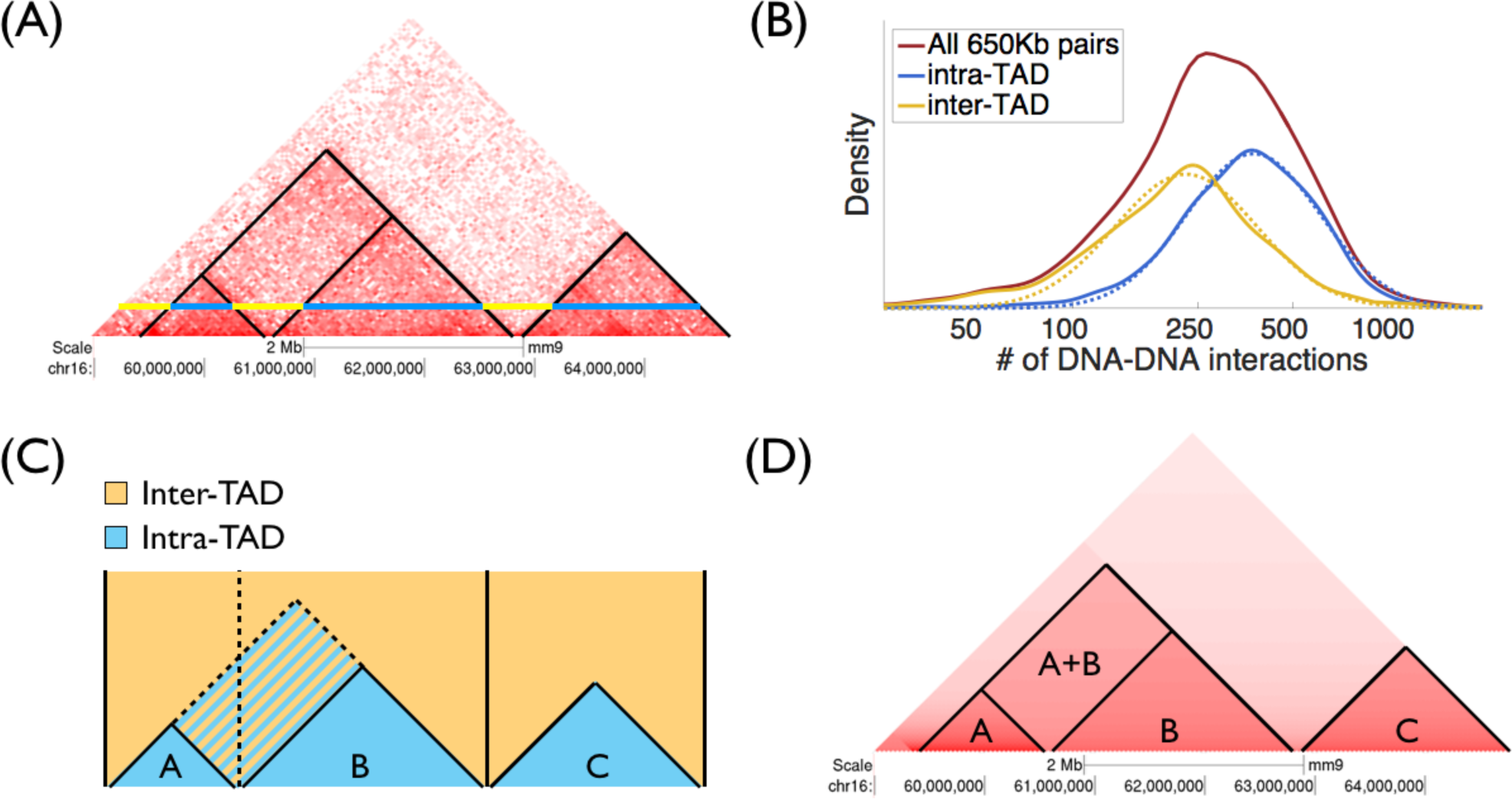
Overview of the PSYCHIC algorithm. **(A)** Example of Hi-C interaction map (rotated in 45°), from mouse cortex (chr16, 59Mb - 64.8Mb) (Dixon et al. 2012). Blue and yellow lines correspond to DNA-DNA pairs, 650Kb apart, within and across domains. **(B)** Histograms show the empirical abundance of DNA-DNA interactions (650Kb apart), located within domains (blue), or across domains (yellow). Dotted lines mark the density function of log-Normal distribution fitted to the empirical data. **(C)** This unified probabilistic mixture model is used to compare the intra- and inter-domain models for each cell in the Hi-C matrix. For example, a proposed segmentation into three domains A-C (delineated by vertical lines), would prefer the intra-TAD model for Hi-C cells within the domains (shown in blue) and the inter-TAD model outside (yellow). An alternative segmentation, where A and B domains are unified would only differ in striped rectangle. Dynamic Programming algorithm identifies the optimal (Viterbi) segmentation of the chromosome into domains. **(D)** PSYCHIC then iteratively merge similar neighboring domains (here, A+B) into a hierarchical structures. Finally, a bi-linear power-law model is used to reconstruct a specific background model for each domain/merge of the Hi-C map, allowing for the identification of overrepresented DNA-DNA pairs, including putative promoter-enhancer interactions.

## Results

### A Unified Probabilistic Mixture Model for Hi-C Data

Hi-C interaction maps often show a clear distinction between two different patterns. Rectangular regions along the diagonal of the Hi-C map correspond to topological domains, and present high intensity of (intra-domain) DNA-DNA interactions. These are often surrounded by regions with fewer (inter-domain) DNA-DNA interactions. Due to symmetry, Hi-C maps are often rotated in 45 degrees, with topological domains shown as isosceles right triangles along the (now horizontal) diagonal of the Hi-C map (Fig. 1A).

We begin by developing a simple two-component probabilistic model, corresponding to the probability of intra- and inter-TAD interactions. In brief, our algorithm analyzes the Hi-C interaction matrix, and infers for every cell (DNA-DNA pair) the Log Probability Ratio (LPR) of these loci occurring within the same topological domain or not. At the following stages we will combine these ratios into a unified score, and use Dynamic Programming to optimally segment each chromosome into domains.

Formally, let ***P_d_(N)*** denote the probability of observing ***N*** Hi-C interactions between two DNA loci ***d*** bases apart. This equals to the sum of the intra-domain and inter-domain sub-models:

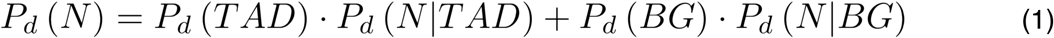

where ***P_d_(N I TAD)*** and ***P_d_(N I BG)*** correspond to the likelihood of observing ***N*** interactions ***d*** bp apart in the intra- and inter-TAD sub-models, respectively. ***P_d_(TAD)*** and ***P_d_(BG)*** correspond to the ***a priori*** probability of observing two loci ***d*** bp apart to be within or outside of the same TAD. For simplicity and robustness, we model N using a log-Normal distribution:

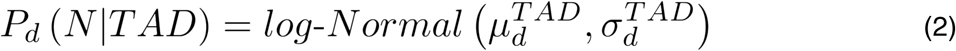

where the log-Normal distribution with mean ***μ*** and standard deviation ***σ*** can be written as:

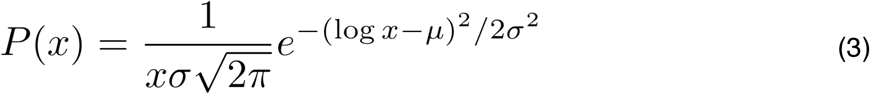

This greatly reduces the number of free parameters, resulting in a compact model ***θ_d_*** with only six parameters for every distant ***d***, including ***μ_d_^TAD^***, ***σ_d_^TAD^***, ***μ_d_^BG^***, and ***σ_d_^BG^*** (mean and standard deviation parameters for intra- and inter-TAD models); and two prior parameters ***P_d_(TAD)*** and ***P_d_(BG),*** while maintaining robust and accurate approximation of the empirical distributions (Figure S1).

For every distance ***d***, we directly estimate the model parameters from annotated Hi-C data. To estimate ***θ_d_***, we rely on an initial (possible noisy) segmentation of the Hi-C map into domains. These could be obtained using various methods, including the directionality index (DI) HMM-based method of Dixon et al (Andersson et al. 2014), or approximated iteratively using the Expectation-Maximization (EM) algorithm (Dempster et al. 1977). Given such annotations, we consider all intra- and inter-TAD pairs and use a maximum likelihood estimation of the mean and the standard deviation parameters. The same approach is used to estimate the prior probabilities, namely which percent of the DNA-DNA interactions of distance ***d*** occur within, or across, topological domains.

### Identification of TAD Boundaries using Log Posterior Ratios

Using the above probabilistic model, we now wish to re-segment the genome into TADs. For this, we propose a score that will integrate information from various distances of DNA-DNA interactions across the entire Hi-C matrix, without being skewed by the significantly higher number of interactions among nearby DNA-DNA pairs.

For this, we define a local score that calculates for every cell in the Hi-C matrix the Log Posterior Ratio (LPR) of the intra- and inter-TAD models. Assuming ***N*** interactions for two DNA loci ***d*** bases apart, we could use Bayes’ law to derive the posterior probability of the intra-TAD model:

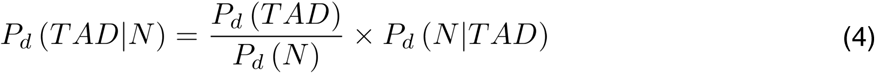

and similarly for the inter-TAD model:

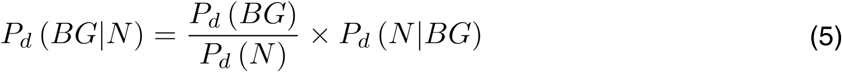

and ***LPR_d_(N)***, the Log Posterior Ratio of the two sub-models could be written as:

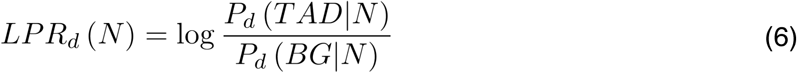

We are now ready to score a segmentation of the genome into domains. First, let us define the probabilistic score for a single topological domain ***t*** from position ***s*** to position ***e***

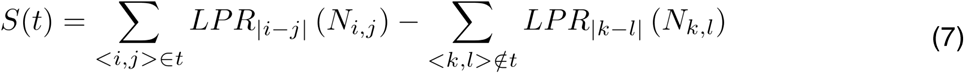

Here, we sum the Log Posterior Ratio for all intra-TAD pairs ***<i,j>*** where ***s ≤ j ≤ i ≤ e***, and subtract the Log Posterior Ratios (or add the log of the inverse ratio) for all inter-TAD pairs of outside TAD ***t***, defined by pairs ***<i,j>*** up to some maximal distance ***h*** (e.g. 4Mb) such that ***s ≤ (I+j)/2 ≤ e.*** These are shown as blue (intra-) and yellow (inter-TAD) regions in Fig 1C. Probabilistically speaking, we allow each Hi-C cell to independently compare its likelihood given each of the two sub-models.

We then define a global score for a segmentation ***C*** of the genome into a set of TADs, by summing over their scores:

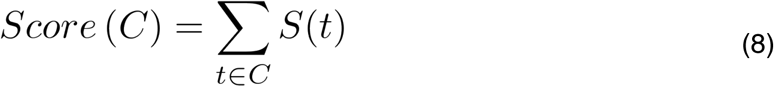

Finally, we find the optimal segmentation of each chromosome into topological domains, with respect to our model. For this, we use a Dynamic Programming algorithm that recursively computes the optimal score of each genomic interval C(i,j) by comparing its score as a one single TAD, or by breaking it at position ***k*** into two distinct regions:

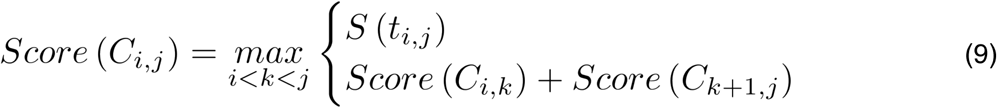

This algorithm allows us to efficiently enumerate over all possible configurations ***{C}*** and identity the optimal segmentation **C**, with respect to the above probabilistic score.

### Hierarchical Model of Topological Domains

So far, we developed a probabilistic framework for modeling Hi-C data within and across topological domains, and presented an efficient algorithm for identifying the optimal segmentation.

For this, our model assumed that all intra-TAD DNA-DNA pairs, located ***d*** bases apart, distribute according to one set of log-Normal parameters, and all inter-TAD pairs use another set.

We now wish to alleviate this assumption, and allow each domain to be modeled by a unique set of parameters. Specifically, we wish to iteratively agglomerative neighboring domains into a hierarchical structure of topological domains. For this, we developed a “merge score” that allows us to examine adjacent domains. A naive scoring system for neighboring TADs would simply quantify their connectivity, by directly counting the number of inter-TAD interactions (Fraser et al. 2015). This score however, might be biased by the size of the two domains, as well as the overall interaction intensity in each of the two domains. Instead, we calculate for each domain the average number of DNA-DNA interactions for any distance, and compare these plots to those of the merged region and inter-TAD regions (Figure S1C). Formally, this translates to finding the optimal ***a*** satisfying:

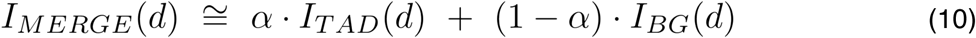

where ***I_MERGE_, I_TAD_***, and ***I_BG_*** denote the average intensities for each ***d*** at the inter-TAD merged area, the two TADs, and at the inter-TAD background model. We do so iteratively, merging the current most similar pair (=highest ***α***), up to a maximal size of 5Mb for the merged structure, thus creating a hierarchical forest-like TAD structure, which corresponds to triangles (TADs) and rectangles (inter-TAD regions).

### TAD-Specific Background Model of Hi-C Data using a Bi-Linear Power-Law Model

Once we have segmented the Hi-C map into hierarchical domains, we wish to model the expected intensity of the Hi-C map. Previous works used a power-law scaling model (Lieberman-Aiden et al. 2009, Mirny 2011, Naumova et al. 2013), to describe ***I*** the number of DNA-DNA interactions as their distance ***Δ*** exponentiated by some coefficient ***a:***

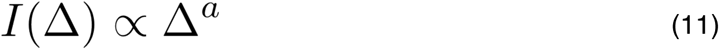

This is often plotted in log-log scale, where the number of interactions (in log scale) scales linearly with the distance (in log scale):

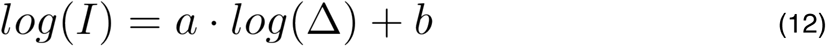

with ***a*** being the power-law coefficient (slope, in log-log plot) and ***b*** is the intersection parameter.

Nonetheless, while we found the power-law model to be generally accurate, it is clear that some domains are characterized with a significantly higher number of interactions than others (Fig 1A), suggesting they would be best described by different power-law parameters (Fig S1C).

We therefore wish to use the hierarchical model of topological domains and construct a local background model of Hi-C intensity, with local parameters (slope ***a_i_*** and intersect ***b_i_***) for each TAD and each inter-TAD merged region (Fig 1D). This will allow us to estimate the expected number of interactions at any distance within every topological domain/merge and quantify the statistical significance over-represented interactions.

Next, we quantified the goodness of fit by each model to the Hi-C data. First, we tested the original segmentation of the genome for the mouse brain Hi-C data (Dixon et al. 2012). For each TAD we estimated the optimal power-law parameters ***a_i_*** and intersect ***b_i_***, resulting with RMSE score of 1.04, an improvement of 9% compared to a random segmentation of the genome (RMSE=1.14. Fig S2). Our segmentation by itself did not yield a better fit (RMSE=1.11), probably due to shorter domains (mean length of 650Kb, compared to 1.5Mb). Following the hierarchical agglomeration of neighboring domains, with additional local background model merge, yielded a much better fit (RMSE=1.02). Finally, we considered a more sophisticated parametric family for modeling Hi-C interaction data. For this, we developed a piecewise linear regression model for modeling the average number of interactions (in log scale) for any distance (in log scale) (Fig S3). This richer power-law model offers a more accurate model (RMSE=0.83), a 20% reduction in the Hi-C fit error compared to the original TAD-specific power-law fit. Put together, the bilinear power-law fit and the hierarchical TAD model allows us to model Hi-C interaction data with high accuracy, thus forming a detailed background model against which we can compare the data and identify over-represented DNA-DNA interactions.

### Gene-Wise Identification of Enriched DNA-DNA Interactions

We now wish to use the hierarchical TAD-specific bi-linear model as background model for Hi-C, and identify over-represented DNA-DNA interactions, that could correspond to promoter-enhancer and other functional interactions in vivo. For this, we aim to compute the “virtual 4C” plot for each promoter, and compare it to the expected number of interactions according to the background model. We consider a large genomic region surrounding each promoter (±1Mb) and search for enriched Hi-C interactions with the promoter. By subtracting the hierarchical Hi-C background model from the actual data, we obtain the “residual” over-representation map. To assign a statistical enrichment score, we model all residual DNA-DNA interactions within this 2Mb window using a Normal distribution, and calculate the Normal p-value of all regions interacting with the promoter, following an FDR correction for multiple hypotheses (Benjamini and Hochberg 1995) (Methods).

We begin by focusing the Foxg1 locus (chr12:50.3Mb-51.2Mb) using Hi-C data from adult mouse cortex (Dixon et al. 2012). Figure 2A shows the residual map for this locus, with two Foxg1 enhancers (hs566 and hs1539) located 550Kb and 750Kb downstream of the gene, with enrichment p-values of 7e-12 and 1e-20, respectively (following FDR correction). These two enhancers were discovered in human by us and others, using ChIP-seq and conservation data (Visel et al. 2007, Visel et al. 2008, Visel et al. 2013). Comparison of our predictions with published ChIP-seq data of H3K27ac, CTCF, and PolII, as well as DNaseI hyper-sensitivity data from the mouse ENCODE project (Mouse ENCODE Consortium et al. 2012), and evolutionary conservation data (Siepel et al. 2005) further identifies the exact location of these Foxg1 enhancers (Figure 2B).

**Figure 2.**
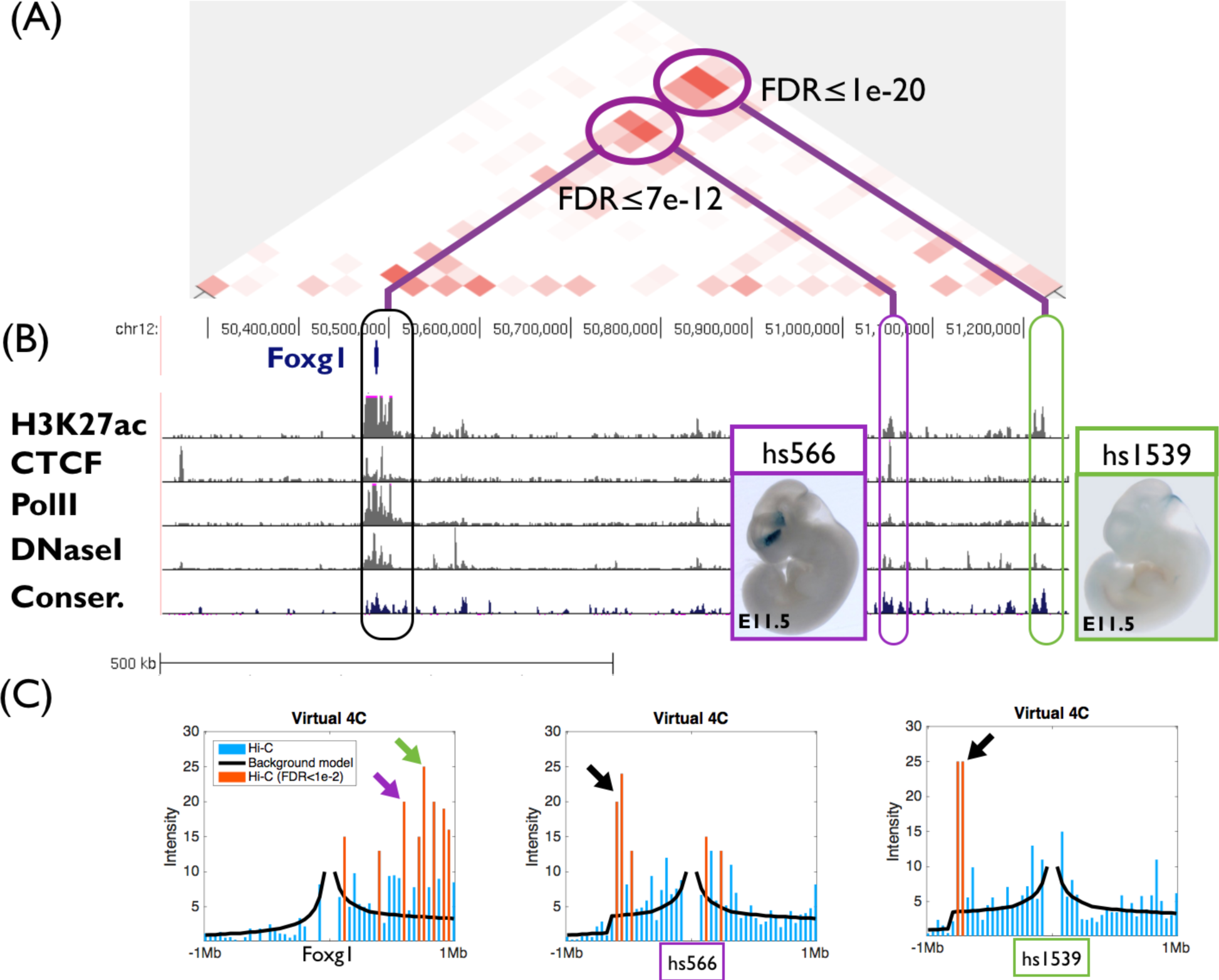
PSYCHIC analysis of the Foxg1 locus in adult mouse cortex Hi-C data (Dixon et al. 2012) identifies two putative enhancer regions, which are enriched with Foxg1. **(A)** Residual map for the Foxg1 locus (chr12:50.3Mb-51.2Mb). These include ChIP-seq marks for active chromatin, and overlap two (human) enhancers validated for brain activity. **(B)** ChIP-seq and conservation data matching active enhancers, within the two putative enhancer regions **(C)** Virtual 4C plots for the Foxg1(left) and the two enhancer loci (hs599, middle; and hs1539 right) loci, comparing Hi-C interaction data against local background model reconstructed by PSYCHIC. Arrows mark significant interactions between Foxg1, hs566 and the hs1539 orthologous regions.

### Genome-Wide Validation of Putative Enhancers

To further test our results on a genome-wide scale, we systematically characterized the chromatin landscape surrounding all predicted enhancers in the mouse cortex (Dixon et al.). For this, we aligned a 4Mb region around each of the 12,278 putative enhancer regions (FDR<1e-2), and compared it to various enhancer-related chromatin marks. These include active enhancer marks (H3K27ac, H3K4me1), promoter marks (H3K4me3, PolII), architectural proteins (CTCF), evolutionary conservation, accessibility, and chromHMM predictions (Siepel et al. 2005, Ernst and Kellis 2012, Mouse ENCODE Consortium et al. 2012, Shen et al. 2012). For all data types, the predicted enhancers were notably enriched compared to their surrounding flanking regions (i.e. regions in 2Mb distance).

Since all predicted enhancers are located no more than 2Mb from known promoters, we wanted to rule this out as a trivial explanation for the observed enrichment. We therefore constructed a similarly sized set of random genomic loci, uniformly sampled around promoters (Fig. 3, red lines). These only show low (15%) enrichment compared to flanking regions.

**Figure 3.**
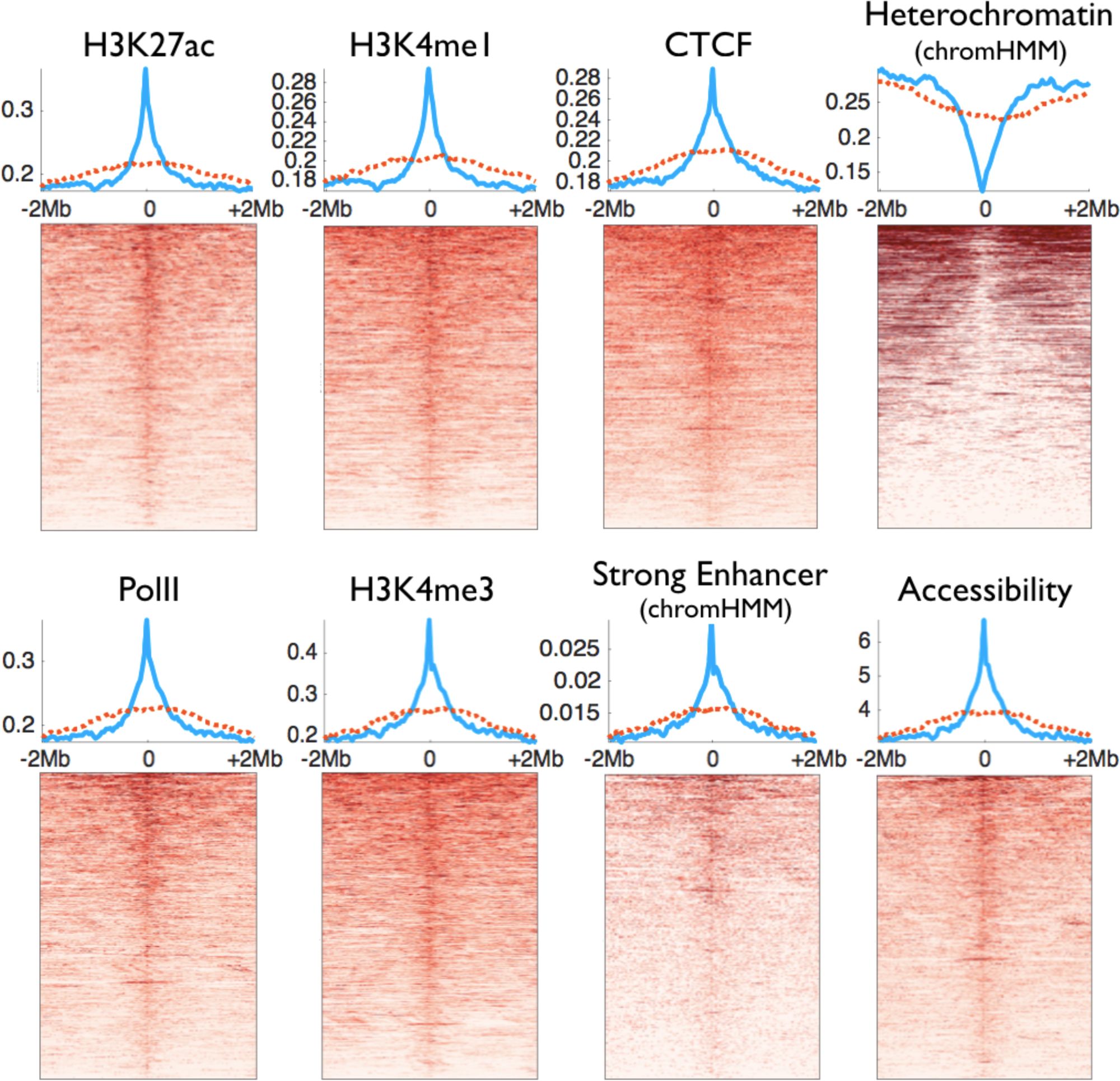
Chromatin marks at 4Mb windows centered around 12,278 putative enhancer regions, predicted using adult mouse cortex Hi-C data (FDR<1e-2) (Dixon et al. 2012). Shown are typical enhancer (H3K27ac, H3K4me1) and promoter (H3K4me3) marks, along with PolII and CTCF ChIP-seq, chromHMM classification, and DNaseI hypersensitivity assays. Blue lines mark the average signal over all predictions. Dotted red lines mark the signal in a random set of genomic loci, sampled in 2Mb windows around promoter.

Notably, most - but not all - putative enhancers show strong enrichment for active chromatin marks. For example, about 70% of the 1e-2 predicted enhancers show increased accessibility compared to their flanking DNA regions (Fig. 3, “Accessibility”). Almost half (46%) of the predicted enhancer regions show enrichment that is greater than one standard deviation compared to their flanking regions (32% > 2SD). For comparison, only 43% of the randomly selected regions show increased accessibility, with only 24% exceeding one standard deviation (15% > 2SD). Similar numbers are obtained for H3K27ac or CTCF.

This suggests that over-represented DNA-DNA interactions (in Hi-C) are not limited to active and accessible regions, and raises the hypothesis that a non-trivial fraction of the putative enhancer regions we have identified are “silent” and inaccessible. A closer examination identified several known enhancers even within those. For example, PSYCHIC identified the ZRS locus as interacting with the *Shh* gene, even in adult mouse cortex (Fig. 4). In the mouse, early developmental *Shh* expression is essential for correct autopod formation, and is regulated in the developing limbs by the distal ZRS enhancer, located ˜1 Mb away (Lettice et al. 2003, Sagai et al. 2005). Our results suggest that ZRS is in close physical proximity to *Shh* even in the adult brain (Fig. 4). This was recently validated by DNA FISH showing ZRS in the proximity of *Shh* throughout a variety of tissues and developmental stages, while not being in active transcription (Williamson et al. 2016).

**Figure 4.**
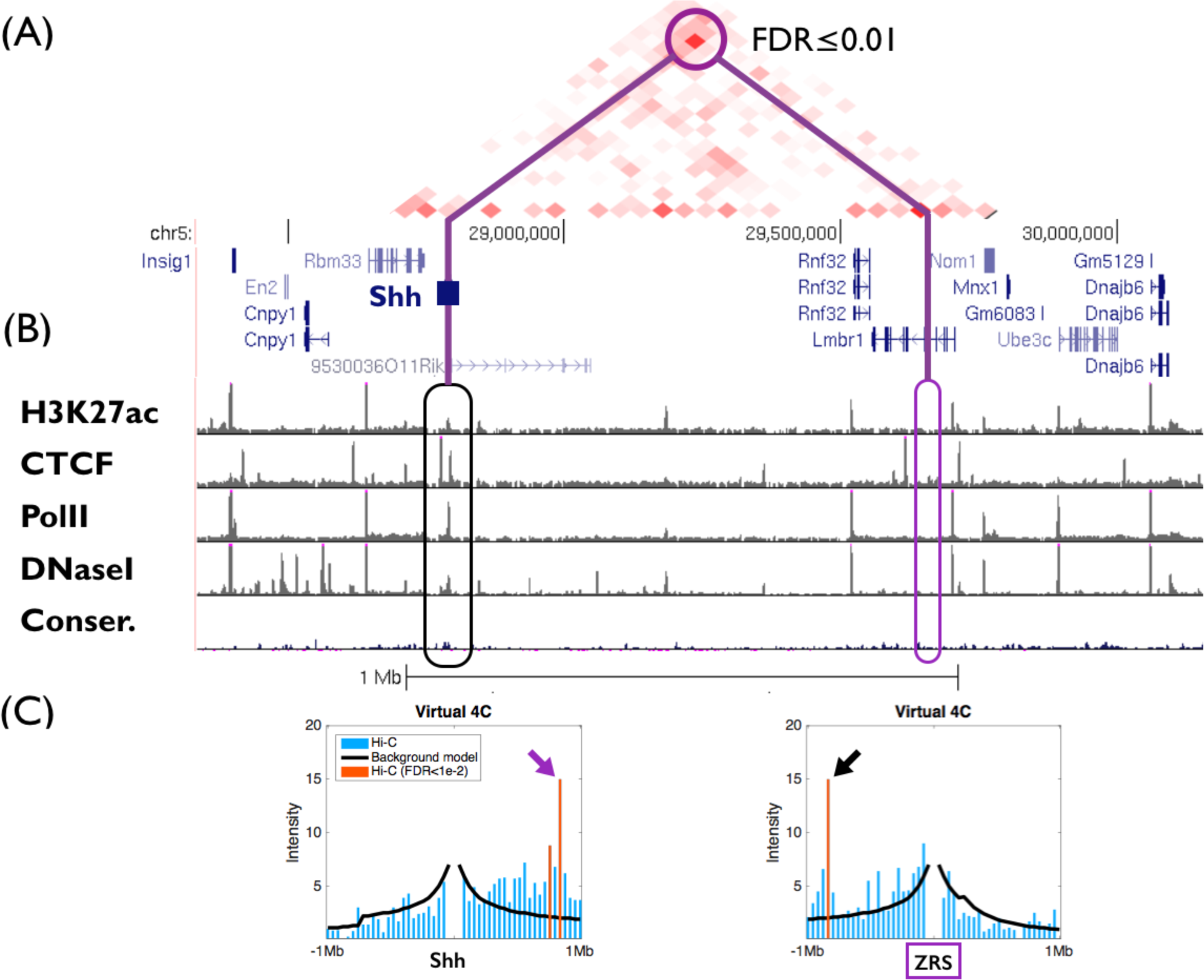
Over-represented promoter-enhancer interactions between *Shh* (in adult mouse cortex) and the limb-specific enhancer ZRS (chr5:28.3Mb-30.2Mb). **(A)** Residual map (of Hi-C data compared to the PSYCHIC hierarchical background fit model) identifies over-represented DNA-DNA interaction between the *Shh* and its limb-specific enhancer ZRS. **(B)** Genome-wide ChIP-seq and accessibility data from adult mouse cortex shows no active enhancer marks for this enhancer, suggesting that ZRS is often interacting with Shh in the brain. **(C)** Virtual 4C plots for the *Shh* (left) and the ZRS (right) loci, comparing Hi-C interactions with the local background model reconstructed by PSYCHIC. Arrows mark significant between *Shh* and ZRS.

### A Comprehensive Catalogue of Human and Mouse Enhancers

To obtain a comprehensive list of putative enhancer regions, we have gathered Hi-C data in 15 conditions and cell types in human and mouse, including mouse cortex and embryonic stem cells (Dixon et al. 2012), mouse embryonic stem cells, neural progenitor cells (NPC), and neurons (Fraser et al. 2015), and mouse B-lymphoblast (CH12LX) cells (Rao et al. 2014), as well as human embryonic stem cells and lung fibroblast IMR-90 cells (Dixon et al. 2012), GM12878 B-lymphoblastoid cells, and HMEC, HUVEC, IMR-90, K562, KBM7, and NHEK cells lines (Rao et al. 2014). Globally, with an enrichment FDR threshold of 0.05, we predicted 320,737 putative enhancers (90,113 in mouse and 230,624 in human) that regulate a total of 27,497 genes (19,016 in mouse and 21,000 in human). A more stringent FDR threshold of 1e-4, yields 123,149 putative enhancer regions (29,732 and 93,417) regulating 22,365 genes (12,603 and 16,919 for mouse and human respectively). These are summarized in Table S1 and on our supplementary webpage www.cs.huji.ac.il/˜tommy/PSYCHIC.

Next, we calculated the distribution over the number of putative enhancers regulating each gene, and compared it to the distribution of randomly selected regions (equivalent to a “random set” of enhancers, chosen with an FDR threshold of 1e-2. See Methods). As shown in Figure S5, for all analyzed Hi-C experiments, we observed a much greater number of genes predicted to be regulated by multiple enhancer regions, compared to the random set. Our results show some genes to be regulated by ten and more enhancers. For example, 443 genes are predicted to have five brain enhancer regions (FDR < 1e-2), compared to only two in the randomized set, or three expected according to a binomial distribution.

Finally, we tested whether the predicted enhancer regions tend to reside within the same TAD as their target genes (Fig. 5). Our analyses suggest that about 88% of predicted enhancer regions (in all 15 analyzed datasets, mouse and human) are indeed within the same domain as their targets, compared to 45% of equally distant random loci. One should note that typically the topological domains called by PSYCHIC are rather short (mean length of 650Kb, compared to ˜1.5Mb for Dixon et al). When considering the inferred hierarchical organization of the genome, we observe the 92% of putative enhancer regions reside within the same TAD or the first level of merging as its target, (Fig. 5, green supplements) compared to 59% at random.

**Figure 5.**
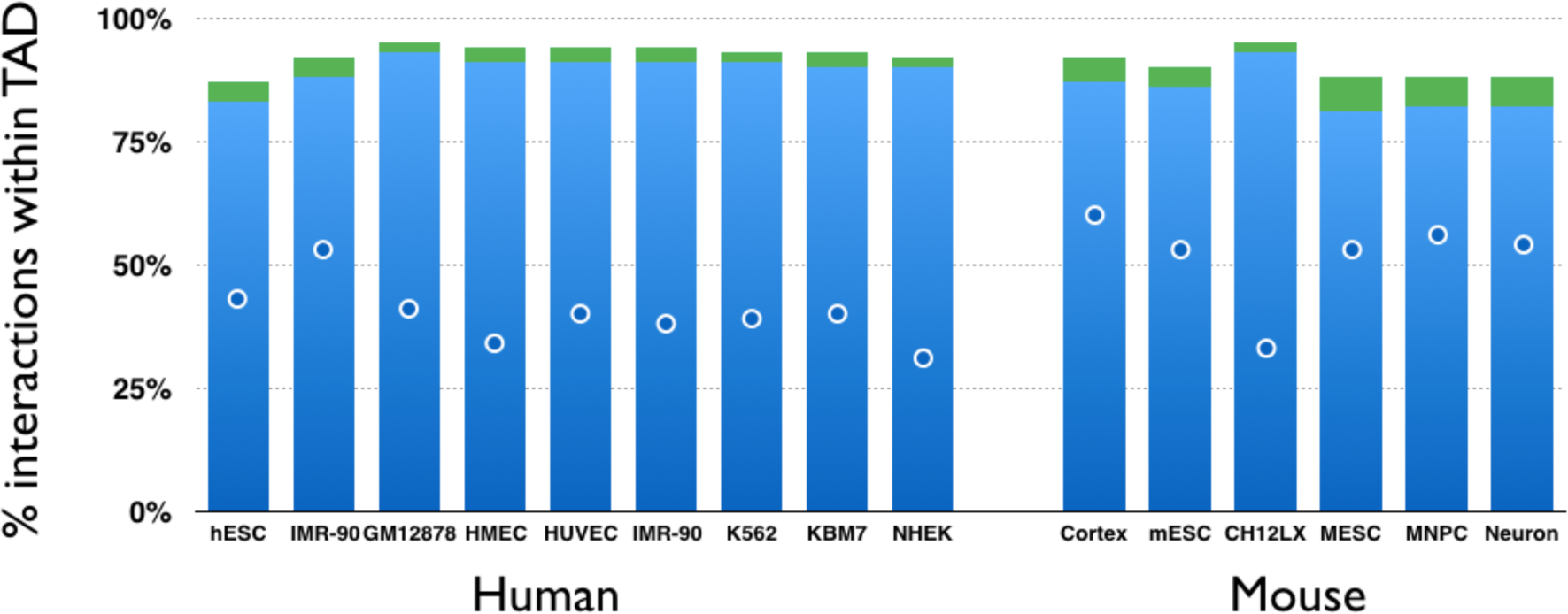
Most putative enhancers reside within the same TAD as their targets. For each of the 15 human and mouse Hi-C experiments we analyzed, the Y-axis shows the percent of predicted DNA-DNA pairs to fall within the same topological domains. Green supplements show the percent of additional pairs falling within 1^st^ level of TAD-TAD hierarchical merges. Blue dots show percent of “random” enhancers residing within the same TAD.

## Discussion

In this work we presented PSYCHIC, a computational model for analyzing Hi-C data to identify enriched DNA-DNA interactions. Using a probabilistic model and efficient algorithms, PSYCHIC identifies the optimal segmentation of chromosomes into topological domains, assembles them into hierarchical structures, and fits a TAD-specific background model for the Hi-C data. By considering a “virtual 4C” plot for every gene, and using the background model for statistical assessments, our algorithm identified 320,737 significant over-represented Enhancer-Promoter interactions in 15 Hi-C experiments in human and mouse.

To segment the genome into TADs, our algorithm uses a probabilistic two-component model that independently computes for every cell in the Hi-C matrix the likelihood ratio between intra-TAD and inter-TAD models. This score assigns similar importance to near and far DNA-DNA interactions, and therefore is less affected by the exponentially higher number of short-range interactions that dominate the Hi-C data, but are mostly invariant of the overall arrangement of the genome in topological domains. In addition, this score is additive and can be easily computed from smaller nested TADs, allowing for a fast and scalable Dynamic Programming algorithm that identifies the optimal segmentation for each chromosome.

For agglomerating individual TADs into hierarchical structures and for the computation of TAD-specific background models, we compute the “interaction spectrum” of each TAD. Specifically, we calculate the average number of Hi-C interactions for DNA-DNA interactions at any distance. While this spectrum was previously modeled by a power-law, our results indicate that replacing the power-law model by a two-segment power-law model greatly improves the model accuracy. Initially, we suspected that this could be due to a mixing effect of two cell populations, each with a different chromosomal organization (and power-law parameters). Alas, this hypothesis cannot hold true, as the sum of two negative power-law functions is always convex, in contrast to the concave behavior of most intensity plots we observe. Instead, these results suggest that the power-law breaking point, typically at 100-300Kb could reflect a transition between two molecular mechanisms used for chromosomal packaging at different hierarchies.

Currently, most available Hi-C data are of rather low resolution varying from 10 to 40Kb. Naturally, this hinders our ability to pinpoint Promoter-Enhancer interactions in high resolution. Nonetheless, various genomic methods for identifying enhancer regions within over-represented DNA-DNA interactions – including ChIP-seq for transcription factors and active histone marks, genomic accessibility, evolutionary conservation or computational sequence-based approaches could all be applied to further analyze putative enhancer regions in higher resolution.

As we showed, both for Foxg1 in the mouse cortex, and later on a genome-wide scale, these putative enhancer regions, defined by over-represented number of Hi-C interactions with promoter regions, typically contain accessible sub-regions that are also enriched for active chromatin marks (H3K27ac, H3K4me1), evolutionary conservation, and are typically often bound by CTCF and PolII. Intriguingly, a closer examination of the data reveals that about a third of the predicted regions are inaccessible and bear no active chromatin marks. These include for example, the ZRS locus that acts as a limb-specific distal enhancer for *Shh*, located nearly ˜1 Mb away. While the ZRS locus shows no accessibility or ChIP peaks in the mouse cortex, and is therefore predicted to be inactive it presents a significant number of interactions with its target gene *Shh*. Indeed, Williamson et al. (2016) recently used FISH and 5C to show that indeed ZRS and *Shh* are located in spatial proximity regardless of their activity.

These results suggest that the 3D structure of the genome may be organized to support regulatory DNA-DNA interactions, rather than merely reflect the set of accessible or active regions of the genome. As more Hi-C is collected and analyzed, we hope to shed light on the causality of gene regulation and genome packaging, as well as the plasticity of genome packaging in general.

Put together, we demonstrated how Hi-C data – typically used to identify TAD boundaries – could be also used to reconstruct a local TAD-specific background model that identifies enriched DNA-DNA interactions, and in particular interactions between enhancers and their target genes.

## Methods

### Piece-wise Linear Regression of log (Intensity) and log (Distance)

We model the Hi-C interaction intensity between two loci as a segmented power-law function of their distance. In log-log scale this is modeled by a two-piece segmented linear regression model. For this, we developed a computational algorithm (implemented in MATLAB) to iterate over the optimal breaking point and estimates the two parameters (intercept and slope) for each segment, while minimizing the squared deviation of the data (in log-log scale). Similarly, a piece-wise linear model was learned for the remaining inter-TAD regions.

### TAD Merges

Neighboring TADs are merged into a hierarchical structure, according to a “merge score” that compares the mean Hi-C intensity per distance within the two underlying TADs, their inter-TAD area, and the null inter-TAD model (represented by ***α*** in Eq. 10). We then iteratively merge the two neighboring TADs whose merge area is the most similar, up to a maximal domain size of 5Mb.

### Random set of enhancers

To obtain a random set of locations along the genome, while maintaining a similar distribution around gene promoters, we considered for each gene all genomic loci up to 1Mb away (on either direction), and selected each with a probability of 1e-2.

### Statistical Enrichment Score

To assign a statistical significance score (p-value) for each putative enhancer (namely, an over-represented interaction between a promoter region and some other locus), we assumed a Normal distribution of the local residual map (i.e. Hi-C minus PSYCHIC background mode) at a 2Mb surrounding the promoter of each gene. We then fitted maximum likelihood estimator for the mean value ***μ_i_*** and its standard deviation ***σ_i_*** and used these statistics to translate the deviation of each Hi-C cell from its background model, into z-scores. Finally, we assigned a p-value for each z-score using a standard Normal cumulative distribution function, and applied a FDR correction for multiple hypothesis (Benjamini and Hochberg 1995).

### Genomic analysis of Putative Enhancers

We used deepTools (Ramírez et al. 2014) to align putative enhancers and generate heatmaps for a 4Mb window surrounding each region, for various genomic data tracks (bigwig files). To estimate the deviation of the putative enhancer location, compared to its surrounding, we estimated the parameters of a Normal distribution based on the two 400Kb regions for each putative enhancer region, located 1.6-2Mb apart on either direction.

### Data availability

PSYCHIC is publicly available via GitHub (ttps://github.com/dhkron/PSYCHIC). A full list of putative enhancer regions, as well as the genes they regulate is available in Supplemental Table S1, and in our supplemental website at www.cs.huji.ac.il/˜tommy/PSYCHIC. Also available in our website are saved UCSC Genome Browser sessions for <u>mouse (mm9)</u> and <u>human (hg19)</u>.

## Acknowledgements

We would like to thank Nir Friedman, Eran Rosenthal, Shira Strauss, and members of the Kaplan lab for helpful discussions and comments.

## Competing interests

The authors declare that they have no competing interests.

**Figure S1.**
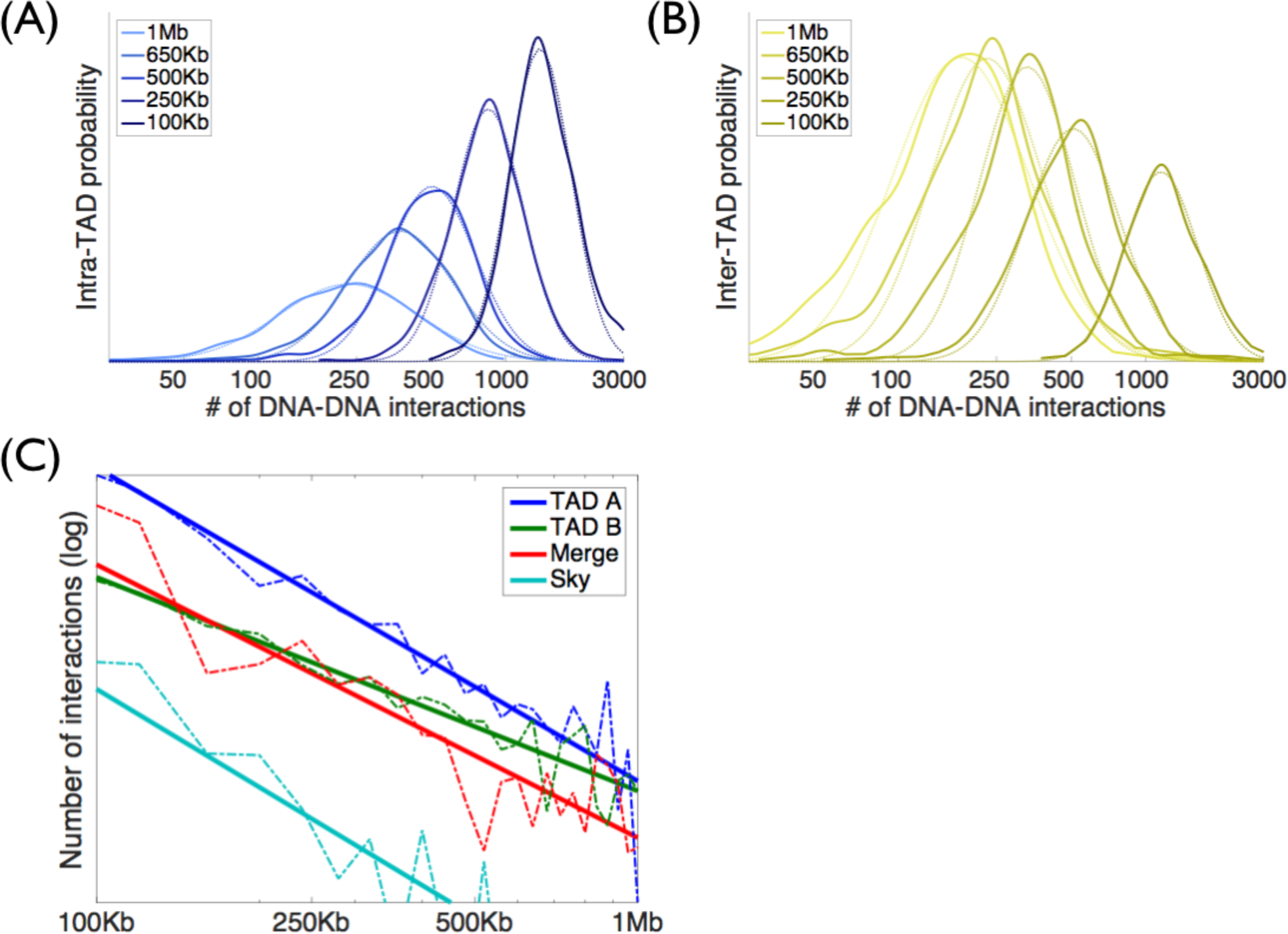
**(A)** Intra-TAD and **(B)** Inter-TAD histograms and matching log-Normal approximations (shown as dotted lines) for DNA-DNA pairs located 100Kb, 250Kb, 500Kb, 650Kb and 1Mb apart. Shown are data from mouse ES cells, chr 11 (Fraser et al. 2015). Distribution were normalized according to their matching *a priori* probabilities, resulting with increased probability for short-range pairs for the intra-TAD models, and long-range pairs for inter-TAD models. **(C)** Power-law distributions for TADs A and B (as in Fig 1), their merged interactions and the inter-TAD background interactions (denoted as “Sky”).

**Figure S2.**
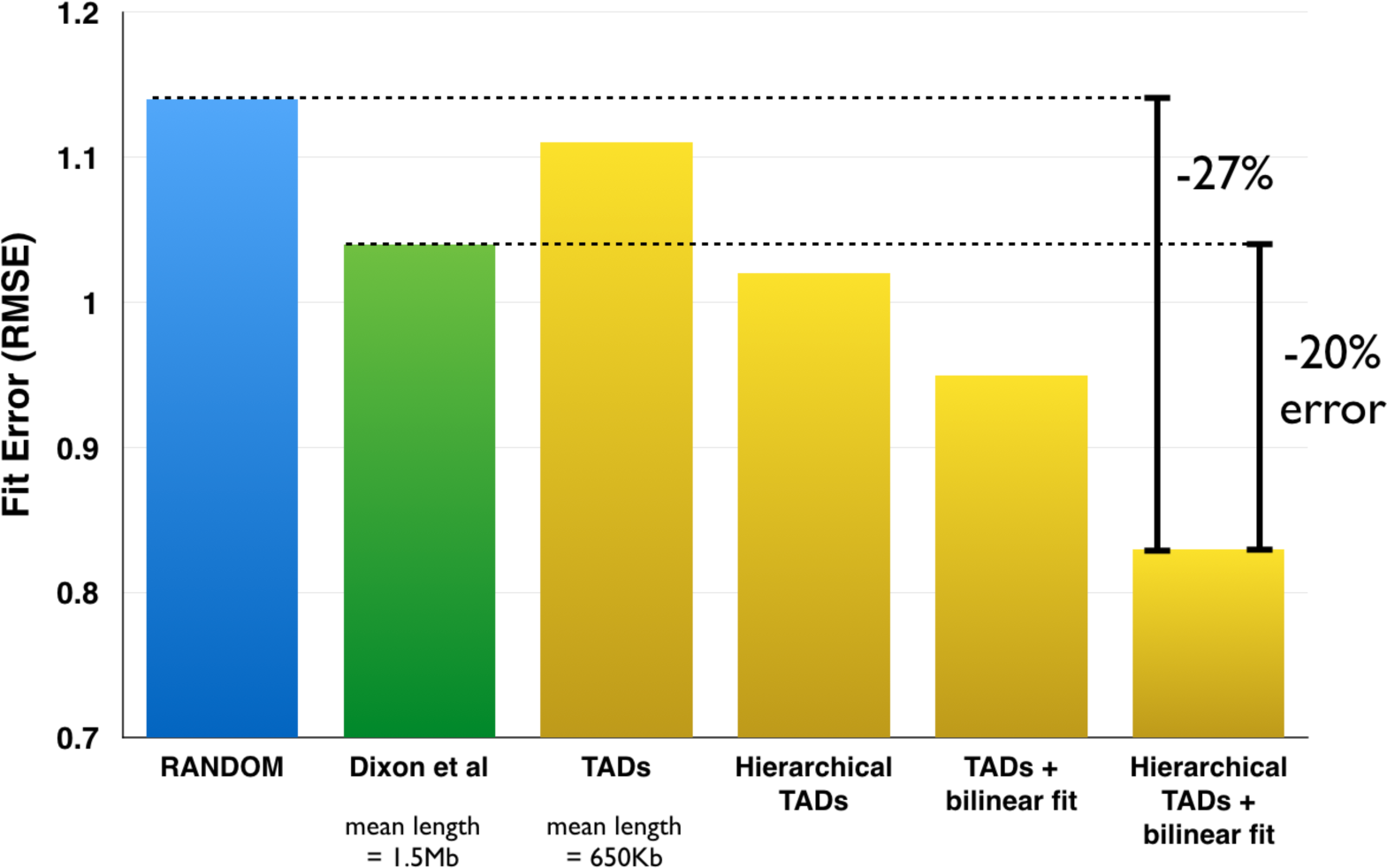
PSYCHIC improves the modeling of Hi-C data by over 20%, compared to similar fit models using the original TAD segmentation by Dixon et al (2012). Here, we compare the root mean squared error (RMSE) of the Hi-C matrix (in log scale) with the reconstructed background model (in log scale).

**Figure S3.**
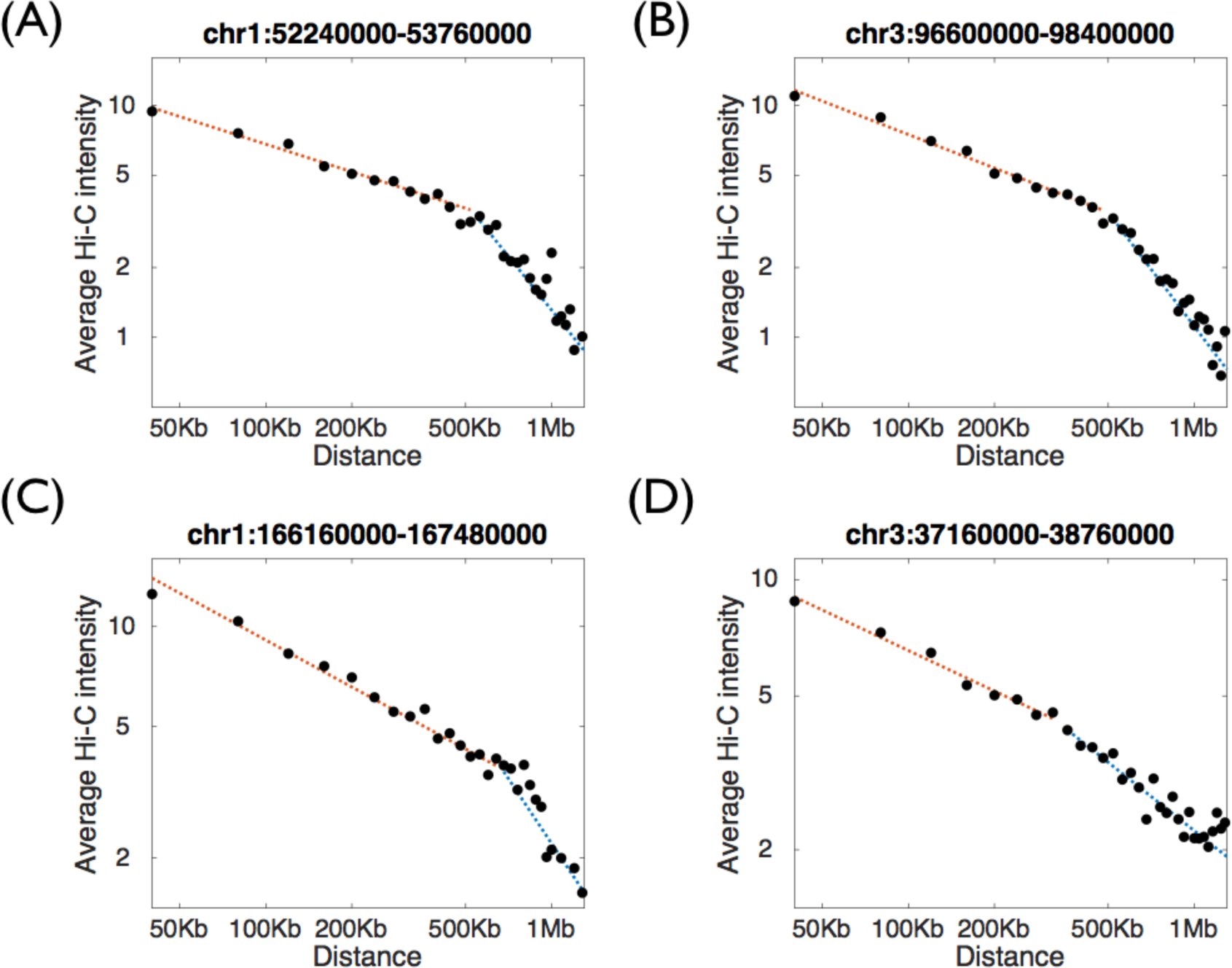
TAD-specific bilinear power-law fit of Hi-C data, for four genomic loci using adult mouse Hi-C data (Dixon et al.). Shown are the average numbers of Hi-C interactions (Y-axis) for each genomic distance between the interacting DNA loci (X-axis). Dotted lines mark the piecewise linear fit.

**Figure S5.**
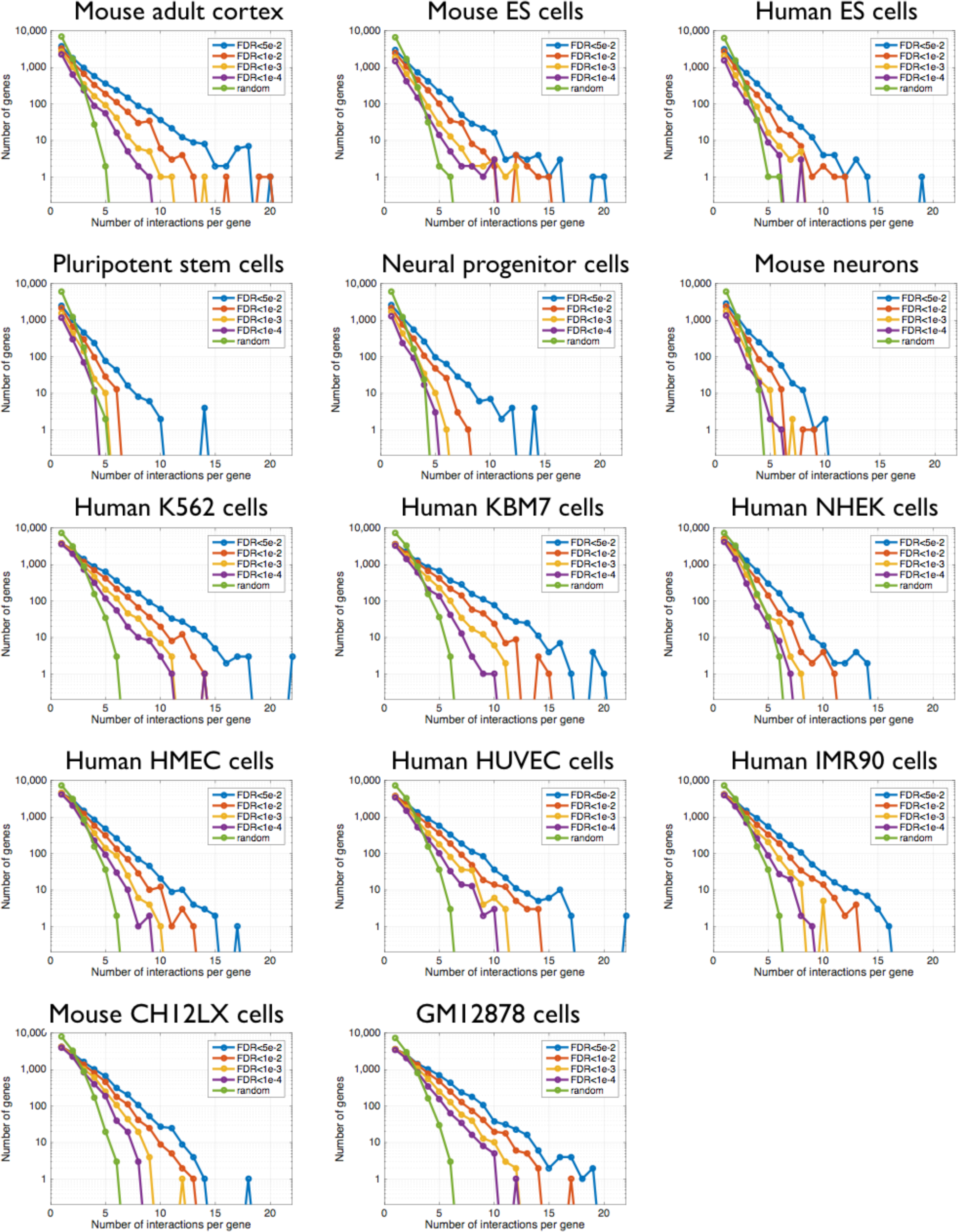
Number of predicted enhancer regions per gene. For each Hi-C dataset, we ran PSYCHIC and predicted putative interactions for each promoter (up to a maximal distance of 1Mb), using several thresholds of statistical enrichment (FDR values of 0.05, 0.01, 1e-3 and 1e-4). Shown are the numbers of genes (Y-axis) predicted to be regulated by X putative enhancer regions (X-axis), compared to a random set of gene-surrounding genomic loci (in green, total size similar to the FDR<1e-2 set of putative enhancers).

## References

1. Achinger-Kawecka J and Clark SJ (2016). Disruption of the 3D cancer genome blueprint. Epigenomics.

2. Adhikari B, Trieu T and Cheng J (2016). Chromosome3D: reconstructing three-dimensional chromosomal structures from Hi-C interaction frequency data using distance geometry simulated annealing. BMC genomics. 17(1): 886.

3. Andersson R, Gebhard C, Miguel-Escalada I, et al. (2014). An atlas of active enhancers across human cell types and tissues. Nature. 507(7493): 455–61.

4. Ay F, Bailey TL and Noble WS (2014). Statistical confidence estimation for Hi-C data reveals regulatory chromatin contacts. Genome Research. 24(6): 999–1011.

5. Benjamini Y and Hochberg Y (1995). Controlling the false discovery rate: a practical and powerful approach to multiple testing. Journal of the royal statistical society. Series B (Methodological). 289–300.

6. Bickmore WA and van Steensel B (2013). Genome architecture: domain organization of interphase chromosomes. Cell. 152(6): 1270–84.

7. Blinka S, Reimer MH, Pulakanti K and Rao S (2016). Super-Enhancers at the Nanog Locus Differentially Regulate Neighboring Pluripotency-Associated Genes. Cell Rep. 17(1): 19–28.

8. Chen J, Hero AO, 3rd and Rajapakse I (2016). Spectral identification of topological domains. Bioinformatics. 32(14): 2151–8.

9. Claussnitzer M, Dankel SN, Kim K-H, et al. (2015). FTO Obesity Variant Circuitry and Adipocyte Browning in Humans. New England Journal of Medicine. 373(10): 895–907.

10. de Laat W and Duboule D (2013). Topology of mammalian developmental enhancers and their regulatory landscapes. Nature. 502(7472): 499–506.

11. Dekker J and Mirny L (2016). The 3D Genome as Moderator of Chromosomal Communication. Cell. 164(6): 1110–21.

12. Demare LE, Leng J, Cotney J, et al. (2013). The genomic landscape of cohesin-associated chromatin interactions. Genome Research.

13. Dempster AP, Laird NM and Rubin DB (1977). Maximum likelihood from incomplete data via the EM algorithm. Journal of the royal statistical society. Series B (Methodological).

14. Dileep V, Ay F, Sima J, et al. (2015). Topologically associating domains and their long-range contacts are established during early G1 coincident with the establishment of the replication-timing program. Genome Research.

15. Dixon JR, Selvaraj S, Yue F, et al. (2012). Topological domains in mammalian genomes identified by analysis of chromatin interactions. Nature. 485(7398): 376–80.

16. Doyle B, Fudenberg G, Imakaev M and Mirny LA (2014). Chromatin loops as allosteric modulators of enhancer-promoter interactions. PLoS Computational Biology. 10(10): e1003867.

17. Ernst J and Kellis M (2012). ChromHMM: automating chromatin-state discovery and characterization. Nature Methods. 9(3): 215–6.

18. Franke M, Ibrahim DM, Andrey G, et al. (2016). Formation of new chromatin domains determines pathogenicity of genomic duplications. Nature.

19. Fraser J, Ferrai C, Chiariello AM, et al. (2015). Hierarchical folding and reorganization of chromosomes are linked to transcriptional changes in cellular differentiation. Mol Syst Biol. 11(12): 852.

20. Fraser P and Bickmore W (2007). Nuclear organization of the genome and the potential for gene regulation. Nature. 447(7143): 413–7.

21. Fudenberg G, Imakaev M, Lu C, et al. (2016). Formation of Chromosomal Domains by Loop Extrusion. Cell reports.

22. Fulco CP, Munschauer M, Anyoha R, et al. (2016). Systematic mapping of functional enhancer-promoter connections with CRISPR interference. Science.

23. Gómez-Marín C, Tena JJ, Acemel RD, et al. (2015). Evolutionary comparison reveals that diverging CTCF sites are signatures of ancestral topological associating domains borders. Proceedings of the National Academy of Sciences. 112(24): 7542–7.

24. Handoko L, Xu H, Li G, et al. (2011). CTCF-mediated functional chromatin interactome in pluripotent cells. Nature Genetics. 43(7): 630–8.

25. Ing-Simmons E, Seitan V, Faure A, et al. (2015). Spatial enhancer clustering and regulation of enhancer-proximal genes by cohesin. Genome Research. gr.184986.114.

26. Jager R, Migliorini G, Henrion M, et al. (2015). Capture Hi-C identifies the chromatin interactome of colorectal cancer risk loci. Nat Commun. 6: 6178.

27. Jin F, Li Y, Dixon JR, et al. (2013). A high-resolution map of the three-dimensional chromatin interactome in human cells. Nature. 503(7475): 290–4.

28. Kieffer-Kwon K-R, Tang Z, Mathe E, et al. (2013). Interactome Maps of Mouse Gene Regulatory Domains Reveal Basic Principles of Transcriptional Regulation. Cell. 155(7): 1507–20.

29. Lajoie BR, Dekker J and Kaplan N (2015). The Hitchhiker’s guide to Hi-C analysis: Practical guidelines. Methods. 72: 65–75.

30. Lettice LA, Heaney SJH, Purdie LA, et al. (2003). A long-range Shh enhancer regulates expression in the developing limb and fin and is associated with preaxial polydactyly. Human Molecular Genetics. 12(14): 1725–35.

31. Lévy-Leduc C, Delattre M, Mary-Huard T and Robin S (2014). Two-dimensional segmentation for analyzing Hi-C data. Bioinformatics. 30(17): i386–92.

32. Lieberman-Aiden E, van Berkum NL, Williams L, et al. (2009). Comprehensive mapping of long-range interactions reveals folding principles of the human genome. Science. 326(5950): 289–93.

33. Lupiáñez DG, Kraft K, Heinrich V, et al. (2015). Disruptions of Topological Chromatin Domains Cause Pathogenic Rewiring of Gene-Enhancer Interactions. Cell.

34. Mifsud B, Tavares-Cadete F, Young AN, et al. (2015). Mapping long-range promoter contacts in human cells with high-resolution capture Hi-C. Nat Genet. 47(6): 598–606.

35. Mirny LA (2011). The fractal globule as a model of chromatin architecture in the cell. Chromosome Res. 19(1): 37–51.

36. Mouse ENCODE Consortium, Stamatoyannopoulos JA, Snyder M, et al. (2012). An encyclopedia of mouse DNA elements (Mouse ENCODE). Genome Biol. 13(8): 418.

37. Naumova N, Imakaev M, Fudenberg G, et al. (2013). Organization of the mitotic chromosome. Science. 342(6161): 948–53.

38. Nichols MH and Corces VG (2015). A CTCF Code for 3D Genome Architecture. Cell. 162(4): 703–5.

39. Nora EP, Lajoie BR, Schulz EG, et al. (2012). Spatial partitioning of the regulatory landscape of the X-inactivation centre. Nature. 485(7398): 381–5.

40. Ong C-T and Corces VG (2014). CTCF: an architectural protein bridging genome topology and function. Nat Reviews Genetics. 15(4): 234–46.

41. Pope BD, Ryba T, Dileep V, et al. (2014). Topologically associating domains are stable units of replication-timing regulation. Nature. 515(7527): 402–5.

42. Ramírez F, Dündar F, Diehl S, Grüning BA and Manke T (2014). deepTools: a flexible platform for exploring deep-sequencing data. Nucleic Acids Research. 42(Web Server issue): W187–91.

43. Rao SSP, Huntley MH, Durand NC, et al. (2014). A 3D map of the human genome at kilobase resolution reveals principles of chromatin looping. Cell. 159(7): 1665–80.

44. Rowley MJ and Corces VG (2016). The three-dimensional genome: principles and roles of long-distance interactions. Curr Opin Cell Biol. 40: 8–14.

45. Ryba T, Hiratani I, Lu J, et al. (2010). Evolutionarily conserved replication timing profiles predict long-range chromatin interactions and distinguish closely related cell types. Genome Research. 20(6): 76170.

46. Sagai T, M H, Y M, M T and T S (2005). Elimination of a long-range cis-regulatory module causes complete loss of limb-specific Shh expression and truncation of the mouse limb. Development. 132(4): 797–803.

47. Seitan VC, Faure AJ, Zhan Y, et al. (2013). Cohesin-based chromatin interactions enable regulated gene expression within preexisting architectural compartments. Genome Research. 23(12): 2066–77.

48. Shen Y, Yue F, McCleary DF, et al. (2012). A map of the cis-regulatory sequences in the mouse genome. Nature. 488(7409): 116–20.

49. Siepel A, Bejerano G, Pedersen JS, et al. (2005). Evolutionarily conserved elements in vertebrate, insect, worm, and yeast genomes. Genome Research. 15(8): 1034–50.

50. Simonis M, Klous P, Splinter E, et al. (2006). Nuclear organization of active and inactive chromatin domains uncovered by chromosome conformation capture-on-chip (4C). Nature Genetics. 38(11): 1348–54.

51. Symmons O, Uslu VV, Tsujimura T, et al. (2014). Functional and topological characteristics of mammalian regulatory domains. Genome Res. 24(3): 390–400.

52. Taberlay PC, Achinger-Kawecka J, Lun ATL, et al. (2016). Three-dimensional disorganization of the cancer genome occurs coincident with long-range genetic and epigenetic alterations. Genome Res. 26(6): 719–31.

53. Tang Z, Luo OJ, Li X, et al. (2015). CTCF-Mediated Human 3D Genome Architecture Reveals Chromatin Topology for Transcription. Cell. 163(7): 1611–27.

54. Van Steensel B and Dekker J (2010). Genomics tools for unraveling chromosome architecture. Nat Biotechnology. 28(10): 1089–95.

55. Vietri Rudan M, Barrington C, Henderson S, et al. (2015). Comparative Hi-C Reveals that CTCF Underlies Evolution of Chromosomal Domain Architecture. Cell reports. 10(8): 1297–309.

56. Visel A, Minovitsky S, Dubchak I and LA. P (2007). VISTA Enhancer Browser—a database of tissue-specific human enhancers. Nucleic Acids Research.

57. Visel A, Prabhakar S, Akiyama JA, et al. (2008). Ultraconservation identifies a small subset of extremely constrained developmental enhancers. Nature Genetics. 40(2): 158–60.

58. Visel A, Rubin EM and Pennacchio LA (2009). Genomic views of distant-acting enhancers. Nature. 461(7261): 199–205.

59. Visel A, Taher L, Girgis H, et al. (2013). A High-Resolution Enhancer Atlas of the Developing Telencephalon. Cell. 152(4): 895–908.

60. Williamson I, Lettice LA, Hill RE and Bickmore WA (2016). Shh and ZRS enhancer co-localisation is specific to the zone of polarizing activity. Development.

61. Xu Z, Zhang G, Wu C, Li Y and Hu M (2016). FastHiC: a fast and accurate algorithm to detect long-range chromosomal interactions from Hi-C data. Bioinformatics.

62. Zhang Y, Wong C-H, Birnbaum RY, et al. (2013). Chromatin connectivity maps reveal dynamic promoter-enhancer long-range associations. Nature.

63. Zuin J, Dixon JR, van der Reijden MIJA, et al. (2014). Cohesin and CTCF differentially affect chromatin architecture and gene expression in human cells. Proceedings of the National Academy of Sciences. 111(3): 996–1001.

